# Increased flexibility of CA3 memory representations following environmental enrichment

**DOI:** 10.1101/2023.10.08.561390

**Authors:** Silvia Ventura, Stephen Duncan, James A. Ainge

## Abstract

Environmental enrichment (EE) improves memory, particularly under conditions of high memory interference^1–6^. Here, we investigate the neural mechanisms facilitating improved memory following EE. Using associative recognition memory tasks that model the automatic and integrative nature of episodic memory, we find that EE-dependent improvements in difficult associative memory discriminations are related to increased adult hippocampal neurogenesis and sparser memory representations across the hippocampus. Additionally, by recording CA3 place cells as enriched and standard-housed rats explored the increasingly distinct shapes of a “morph” box, we report for the first time that EE changes the way CA3 place cells discriminate similar contexts. CA3 place cells of enriched rats show greater spatial tuning, increased firing rates, and enhanced remapping to contextual changes. These findings point to more precise and flexible CA3 memory representations in enriched rats. Increased spatial tuning and flexibility might support the well-established EE-dependent improvements in fine memory discrimination.

**Highlights:**

- EE improves discrimination of similar associative memories
- EE-dependent memory improvements are related to increased adult neurogenesis
- EE results in sparser hippocampal activity
- EE increases spatial tuning, firing and remapping of CA3 place cells

## Introduction

The ability to discriminate across similar past experiences is a fundamental feature of episodic memory. The hippocampus is thought to support this ability via pattern separation, or the encoding of similar events using dissimilar memory representations^7^. Spatially tuned pyramidal cells in the hippocampus (place cells) contribute to pattern separation by changing their firing patterns in response to spatial and contextual alterations to the environment^8–14^. This process, referred to as remapping^8^, results in events experienced in similar contexts being encoded as different memories. Two types of remapping have been described. Global remapping occurs when distinct populations of place cells are recruited to represent different spatial contexts^8,11,15^. Rate remapping occurs when the same populations of place cells encode different contexts by changing their firing rate while maintaining their firing location^12,15–17^. Both forms of remapping provide a mechanism to reduce the chance of memory interference.

Adult hippocampal neurogenesis has been consistently implicated in pattern separation^18–21^. Adult-born granule cells (abGCs) contribute to pattern separation by driving inhibition in downstream hippocampal regions^19,22–26^, thus leading to sparser, largely non-overlapping hippocampal representations of similar events. Consistent with this, environmental enrichment (EE), which upregulates adult hippocampal neurogenesis^27^, improves rodent performance in several memory tasks involving high memory interference^1–6^. However, pattern separation tests have so far failed to model many of the key features of episodic memory such as automatic encoding and integration of episodic memory features. This is important as failure to model episodic memory effectively limits our ability to apply these findings in a clinical setting which could account for failures to translate basic memory research into treatments for Alzheimer’s disease.

Improvements in pattern separation are associated with functional changes within the hippocampal network, including increased sparseness of neuronal activity and enhanced global remapping of CA1 place cells as rats move between distinct environments^28^. However, whether EE improves memory discrimination by promoting remapping in CA3 place cells that receive input directly from abGCs in DG^22,29^ has not been investigated. Interestingly, recent evidence suggests that CA3, particularly its proximal portion (closer to the DG), is involved in pattern separation^30–32^, with CA3 place cells rate remapping in response to small changes to the environment^12,16,33^. Consistently, stimulation of immature abGCs, which are involved in pattern separation^18–21^, results in increased place cell remapping in proximal CA3^34^. Here, we investigated whether EE modulates place cell activity and remapping in CA3.

We report that EE enhances discrimination of similar contextual memories during an associative recognition memory task, which models episodic-like memory in rodents. These improvements were associated with increased levels of adult hippocampal neurogenesis and sparseness of activity across the hippocampus. Additionally, EE enhanced spatial tuning, firing rates and remapping of CA3 place cells in response to small changes to the environment. Together, these findings point to increased adult hippocampal neurogenesis, enhanced sparseness of hippocampal activity and CA3 place cell remapping as potential mechanisms for enrichment-dependent memory improvements.

## Results

### Improved discrimination of similar associative memories and increased adult hippocampal neurogenesis following EE

To investigate whether EE improves discrimination of similar episodic-like memories, we tested enriched and standard-housed rats on the Object-Context (OC) and Object-Place-Context (OPC) recognition memory tasks, which test spontaneous memory for associations of objects with places and/or contexts. The effect of EE on rodent performance in these tasks has not been investigated before. To manipulate the overlap in contextual information, the animals were tested on two versions of the tasks (Figures 1A,B). In the similar condition, the contexts used were the square and circle version of a “morph” box which shared the same walls and floor. In the dissimilar condition, the contexts were distinct boxes with different walls and floors making them easier to discriminate.

**Figure 1:**
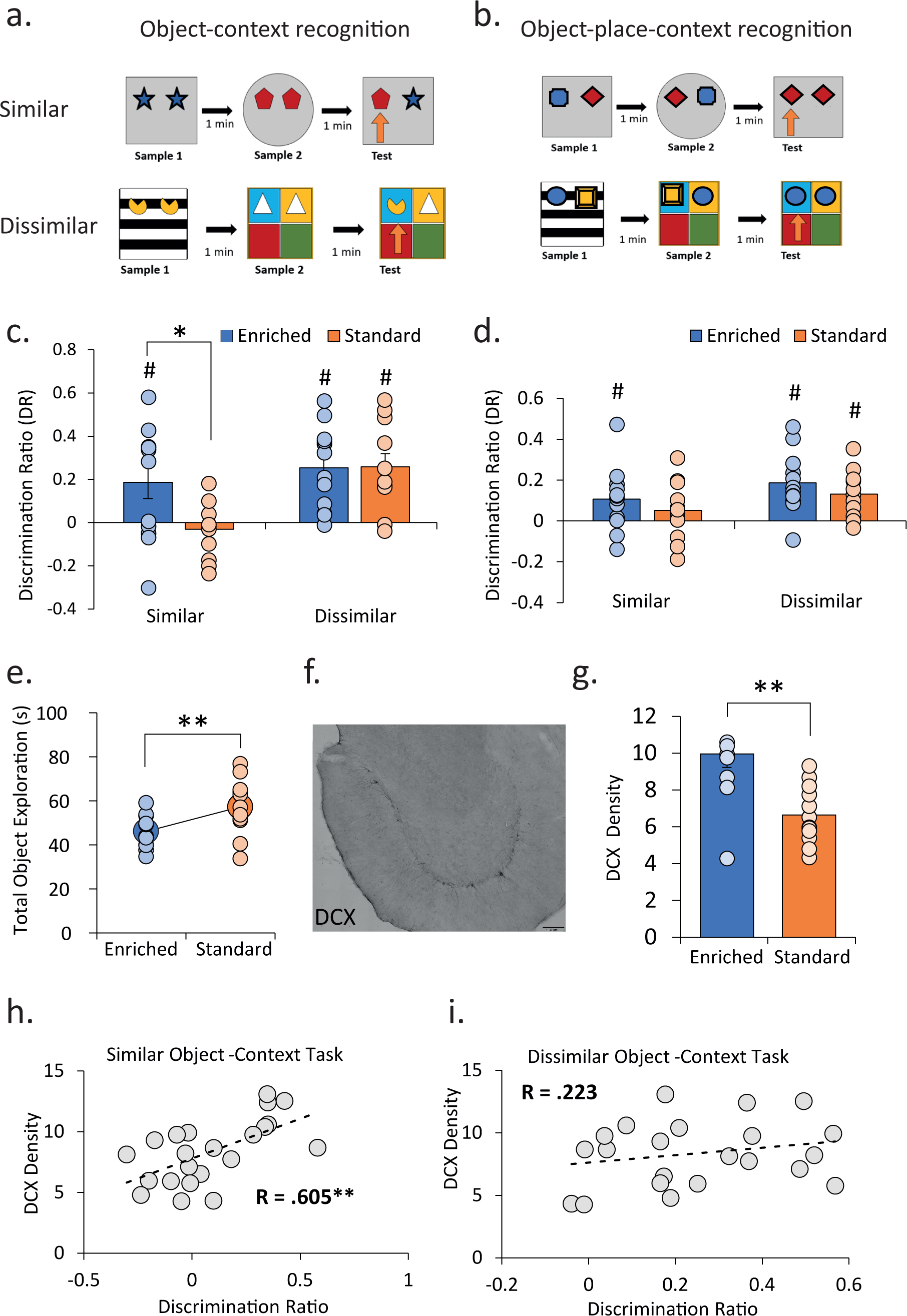
Improved Discrimination of Similar Associative Memories and Increased Adult Hippocampal Neurogenesis following EE. (A and B) Schematic of the object-context (OC) (A) and of the object-place-context (OPC) recognition memory tasks (Top, similar. Bottom, dissimilar). Orange arrow indicates object in the novel configuration. (C and D) Discrimination ratios for the OC (C) and OPC (D) tasks. Bars indicate average discrimination ratios for enriched and standard rats in the similar and dissimilar versions of the OC and OPC tasks. Dots indicate values single animals. Error bars are SEM. *p<0.05. (E) Line graph shows the average total object exploration times during encoding for the enriched and standard groups across the OC and OPC tasks. Dots indicate values for single animals. **p<0.01. (F) Horizontal brain sections showing expression of Doublecortin (DCX) in the dentate gyrus Scale bars represent 37 μm. (G) Bars indicate average density of DCX + cells in the DG of the enriched and standard-housed rats. Dots indicate values for single animals. Error bars are SEM. ***p<0.001. (H and I) Scatterplots of the density of DCX+ cells plotted against the discrimination ratios for the Similar OC (H) and Dissimilar OC (I) tasks. Line of best fit (dashed black) is shown for each plot. **= p<0.01.

Rats raised in an enriched environment for four months (EE, N=12) performed significantly better than standard-housed rats (ST, N=12) in the similar (*F*_(1,21)_ = 6.092, *p* = 0.022, η_p_^2^ =. 22), but not dissimilar (*F*_(1,21)_ = .002, *p*=.962, η_p_^2^=.000), OC memory task (Figure 1C), pointing to a selective enrichment-dependent improvement in fine associative memory discrimination. Critically, while both groups performed above chance level in the dissimilar version of the OC task, showing successful discrimination of distinct contextual memories, only the enriched rats performed above chance level when the contexts used were more similar. In the OPC task, only the enriched rats showed memory for the similar object-place-context memories. However, there was no significant difference in performance across the groups in either the similar or the dissimilar version of the OPC task (Similar: *F*_(1,21)_ = .762, *p*=.393, η_p_^2^ =.035; Dissimilar: *F*_(1,21)_ =. 958, *p*=.339, η_p_^2^=.044) (Figure 1D). Interestingly, the enriched rats spent less time exploring the objects both in the similar and in the dissimilar versions of the tasks during encoding (Figure 1E). Together, these findings suggest that EE leads to improved memory discrimination of similar the contextual features of episodic-like memories, and more efficient encoding of novel associations in episodic memory.

Next, we examined adult hippocampal neurogenesis in the dorsal hippocampus (Figure 1F) by measuring density of DCX positive cells. Enriched rats had increased DCX positive cell density in the dentate gyrus region compared to the standard-housed rats (*F*_(1,21)_ = 15.10, *p*<0.001, η_p_^2^=.42) (Figure 1G). Additionally, there was a positive correlation between levels of adult hippocampal neurogenesis and performance in the similar, but not dissimilar, OC task (Similar: *r*_(22)_ = .605, *p* = .003; Dissimilar: *r*_(22)_ = .223, *p*=.852; Figure 1H,I). There was, however, no correlation between performance in either the similar or the dissimilar OPC task and levels of adult hippocampal neurogenesis (S1C, S1D). These findings show that EE leads to greater levels of adult hippocampal neurogenesis, and improved discrimination of similar spontaneous context-dependent associative memories.

### Increased sparsity of activity in the hippocampus of enriched rats

Previous studies point to EE resulting in sparser hippocampal memory representations ^28^. This is consistent with immature abGCs, which are upregulated by EE^27^, driving inhibition across the hippocampus ^19,22–26^. Increased hippocampal inhibition is thought to promote pattern separation of similar events and reduce interference^20^. Next, we assessed whether EE resulted in increased sparsity in the hippocampus as rats explored novel and familiar object-context associations using *c*-*Fos* as a marker for neural activity (Figure 2A). Rats explored two copies of one object within one of the similar environmental contexts (square or circle) over 2 days. On day 3, half of the rats experienced the same objects within the same context (familiar object-context condition) or within the opposite similar context (novel object-context condition) (Figure 2B). This allowed us to assess differential effects of EE on hippocampal activity in response to novel and familiar object-context configurations.

**Figure 2:**
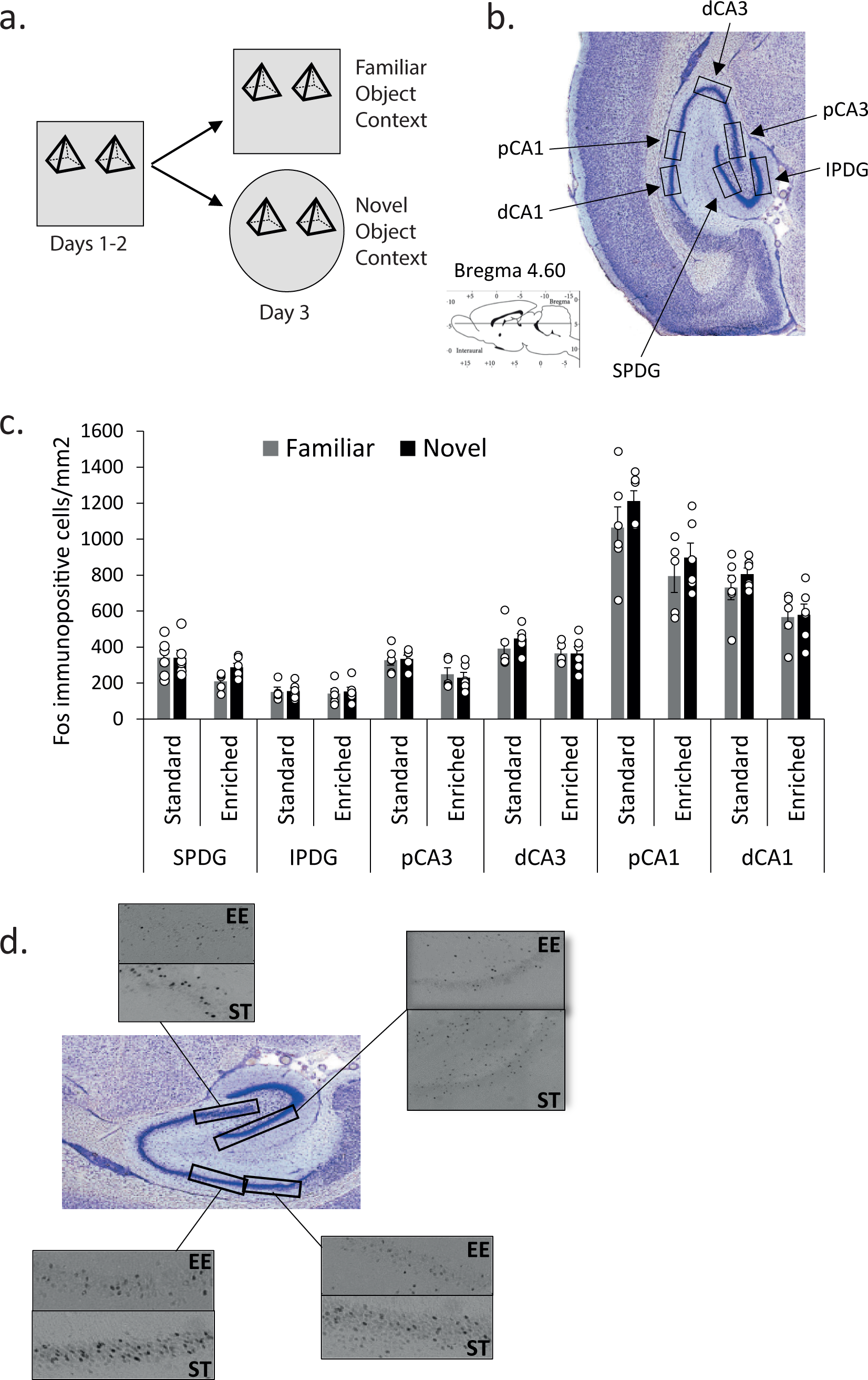
Increased Sparsity of Activity in the Hippocampus of Enriched Rats. **(A)** Schematic of the novel and familiar conditions of the *c-fos* experiment. **(B)** Horizontal section stained with NeuN. Black boxes indicate regions of interest (ROIs). SPDG: suprapyramidal dentate gyrus, IPDG: infrapyramidal dentate gyrus, pCA3: proximal CA3, dCA3: distal CA3, pCA1: proximal CA1, dCA1: distal CA1. **(C)** Bar graph showing average *c-fos* cell densities in the different subregions of hippocampus. Error bars indicate SEM. Data are presented for the enriched and standard-housed rats in the novel and familiar conditions. **= p<0.01. **(D)** Representative images of horizonal sections showing *c-fos* expression from the SPDG, pCA3, pCA1 and dCA1 of the enriched and standard-housed rats.

We found that EE led to reduced activation across the hippocampus in response to both novel and familiar object-context configurations (*F*_(1,19)_ =11.927, *p* = 0.003, η_p_^2^ =.386). There was also a significant interaction of hippocampal subregion and enrichment group (*F*_(5,95)_ =7.188, *p* < 0.001, η_p_^2^ =.274). Reduced activation in enriched rats was particularly evident in the suprapyramidal dentate gyrus (SPDG) (*p* = 0.016), proximal CA3 (*p*= 0.005), and proximal and distal CA1 (*p* = 0.004; *p* = 0.003; Figure 2D). Together, these findings show that EE leads to sparser memory representations of similar novel and familiar object-context associations in the hippocampus.

### Increased spatial tuning and firing rates of CA3 place cells of enriched rats

To investigate the effect of EE on place cell activity, we implanted tetrodes in CA3 of enriched (N=5) and standard-housed rats (N=5) (Figure 3A). We focused on CA3 as place cells in this subregion receive direct input from abGCs^22,29^, and yet the effect of EE on CA3 place cells is unknown. Place cells were recorded as the rats explored the distinct versions of a “morph” box that could be shaped as a square, a circle or as four intermediate shapes^16,17,33^ (Figure 3B, Figure S2B). Each session began with exploration of the square or circle, followed by exploration of the intermediate shapes (1:7, 2:6, 3:5, 4:4 or 4:4, 3:5, 2:6, 1:7), the opposite shape (circle or square) and a repetition of the first shape. Analysis of CA3 place fields from the square box (10 rats; 5 enriched, 5 standard, Table 1) showed that in the enriched rats CA3 place cells had smaller place fields (Figure 3C) (*H*_(1)_ = 6.948, p=0.008) and carried greater spatial information (Figure 3D) (*H*_(1)_ = 19.115, p<0.001) compared to CA3 place cells in the standard-housed rats. EE also resulted in reduced sparseness of firing (Figure 3E) (*H*_(1)_ =19.546, p<0.001), enhanced spatial selectivity (Figure 3F) (*H*_(1)_ 16.821, p<0.001), and increased firing rates (Average In-Field Rate - *H*_(1)_ = 13.700, p<0.001; Peak Rate - *H*_(1)_ =12.381, p<0.001) (Figure 3G-H) (Figure 3I for example cells). Together, these findings point to increased spatial tuning of CA3 place cells in the enriched rats. This is accompanied by an increase in gain, signalled by the upregulation of firing rates.

**Figure 3:**
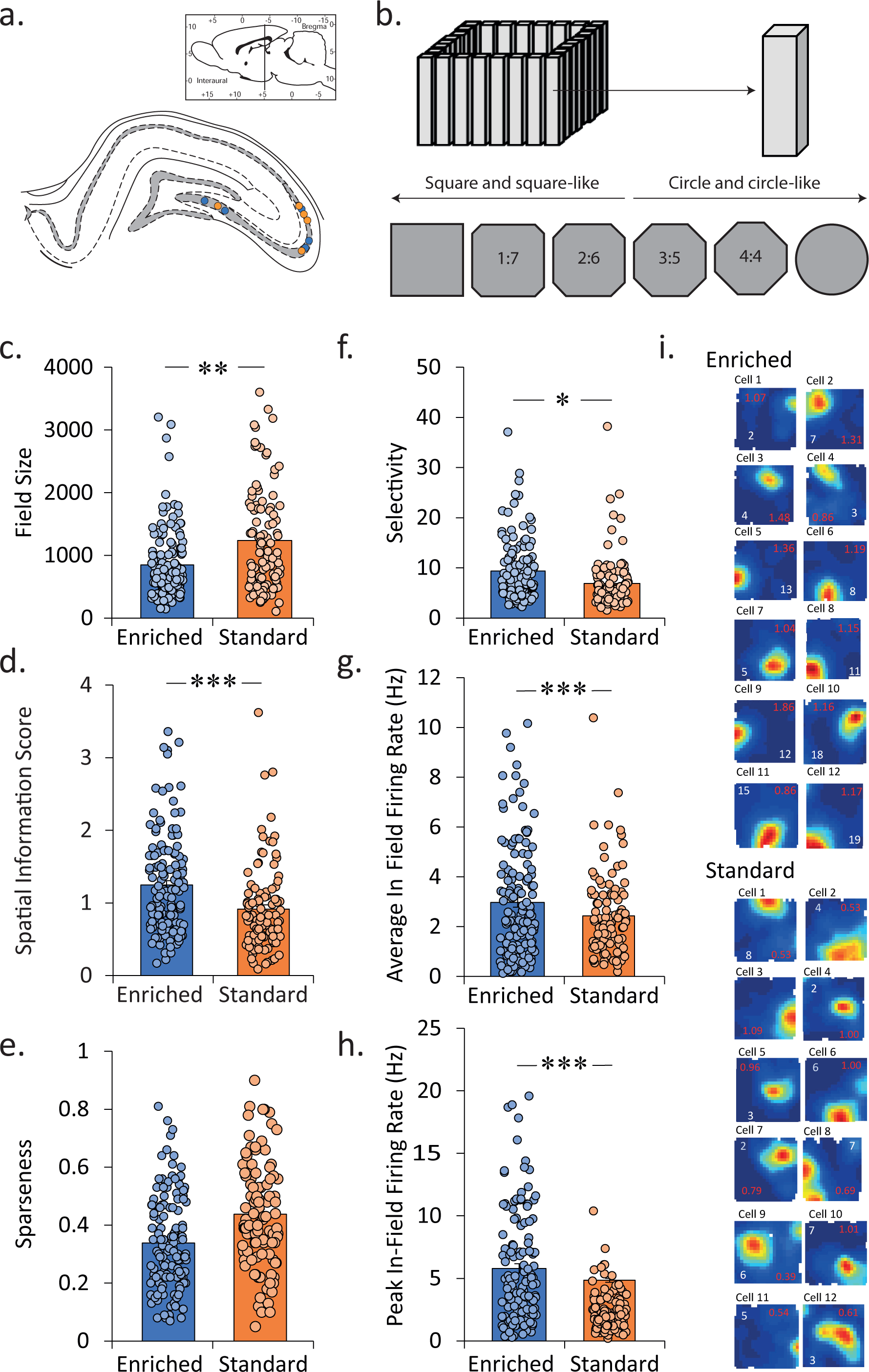
Increased Spatial Tuning and Firing Rates of CA3 Place Cells of Enriched Rats. **(A)**Schematic of tetrode tracks from the enriched (blue) and standard-housed (orange) rats. **(B)** Schematic of the morph box shaped as a square (Top) and of the morph environments (Bottom). **(C-H)** Bar graphs comparing field size (C), spatial information score (D), sparseness (E), selectivity (F), average in-field firing rate (G) and peak in-field firing rate (Hz) (H) across the enriched and the standard-housed rats. Dots indicates values for single place fields. *p<0.05, **p<0.01, ***p<0.001. Error bars are SEM. **(I)** Rate maps from representative cells recorded from the enriched (Top) and the standard (Bottom) groups. Warm colours = high firing rates, cool colours = low firing rates or no firing. Peak firing rates for each rate map are shown in white, spatial information scores are in red.

### Enhanced place cell remapping and increased subset switching in CA3 place cells of enriched rats

Next, we sought to assess whether EE modulates CA3 place cell remapping. Previous studies using the morph environment reported a context-dependent firing rates redistribution (rate remapping) in CA3 place cells as rats explore the shapes of the morph box when they were trained with the square and circle shapes in the same location^16,17,33^. Here, we assessed whether EE affects CA3 place cell remapping in response to small changes to the environment in the morph box. To investigate the effects of EE on rate remapping, we analysed all cells that had a field in all the shapes included in the recording session (full sequence) (Figure 4A, S2C, Table 2). CA3 place cells from the enriched rats showed increased rate remapping across the sequence (*F*_(1,1203)_=7.653, *p* = 0.006) (Figure 4B,C). EE led to increased rate remapping across the square and the circle shapes, as well as across the square and the most circle-like shapes. Interestingly, CA3 place cells in the enriched rats also showed increased rate remapping in response to the delayed exploration of the initial shape. These findings point to EE resulting in increased flexibility in the CA3 spatial map, as CA3 place cells in the enriched rats showed greater ability to change their firing patterns in response to contextual alterations, as well as to a delayed exploration of a familiar environment. Increased rate remapping was reported across all enriched rats (Figure S2C). Interestingly, in the same pool of cells, the spatial correlations between rate maps across recordings from distinct shapes within the session remained high (Figure S2D) and there was no difference in spatial correlations across groups. This shows that CA3 place cells encode fine changes in context (both spatial/geometric and temporal) using rate rather than global remapping. We next aimed to assess whether EE influenced global remapping across the morph sequence in all recorded place cells, including those with a place field in at least one shape in the session in which they were recorded (Table 3) Consistent with the findings reported above, there was no difference in spatial correlation values for the place cells across the groups (Figure 4D) (*F*_(1,346.18)_=1.302, *p*=.255). Together, these findings suggest that EE does not alter global remapping of CA3 place fields.

**Figure 4:**
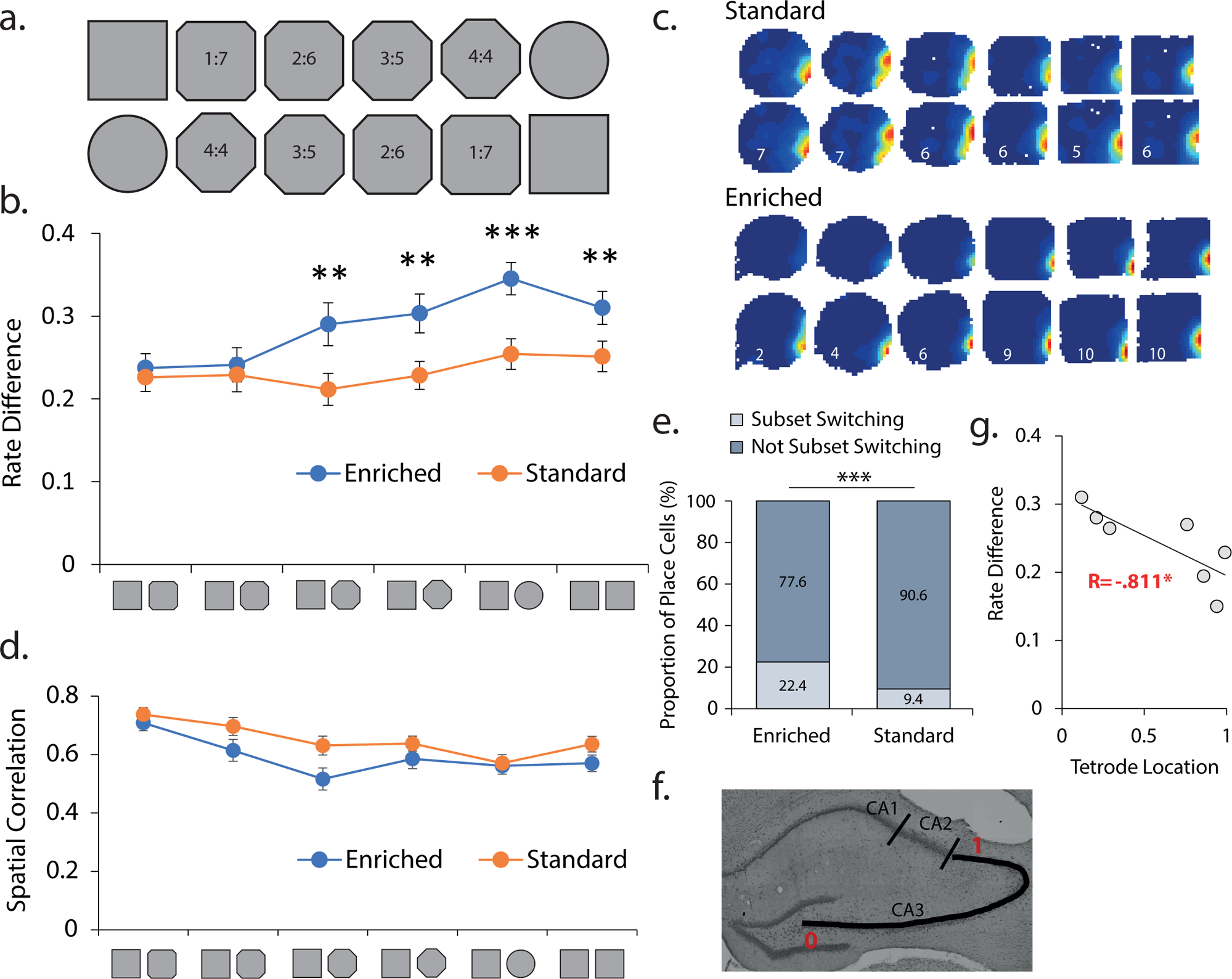
Enhanced Place Cell Remapping and Increased Subset Switching in CA3 Place Cells of Enriched Rats. **(A)** Schematic of the full sequence. **(B)** Line graph showing the average rate remapping scores across the first shape in the sequence and each subsequence shape for cells with a field in all the environments in the recording session **p<0.01, ***p<0.001.**(C)** Rate maps for example place cells recorded across the full sequence from standard (Top) and enriched (Bottom) rats. For each cell, the maps are shown scaled to the maximum firing rates across all environments (Top) and to the cell’s own maximum firing rate within a session (Bottom). **(D)** Line graph showing the average spatial correlation scores across the first shape in the sequence and each subsequence shape for cells with a field in at least one of the environments in the recording session. **(E)** Stacked bar plot comparing the proportion of place cells showing subset switching (turning on an off) across the recording sequence. ***p<0.001.**(F)** Tetrode location across the proximodistal axis of CA3. 0 = most proximal, 1 = most distal. **(G)** Scatterplot of the tetrode location plotted against the rate remapping values across the full sequence for enriched and standard-housed rats. *p<0.05. Line of best fit (dashed black) is shown for each plot.

Place cells might also support discrimination of distinct environments by switching on and off as the environment changes, i.e., subset switching^8,35^. Subset switching is thought to result in distinct cell population encoding distinct environments, thus reducing memory interference. Long-term EE results in increased subset switching in CA1 as rats explore distinct spatial environments^28^. We compared the percentage of place cells active in at least one, but not all, environments in the enriched and standard groups. Critically, a greater percentage of place cells exhibited subset switching in the enriched rats (44/196 place cells; 22.4%) compared to the standard rats (15/160 place cells; 9.4%) (χ^2^_(1)_= 10.890, p<0.001) (Figure 4E). This was the case even when conducting the analysis on a per animal basis (t_(8)_=2.541, p = 0.035, d’= 8.360) (Figure S3A). These findings point to more specific encoding of environments following EE.

Finally, as place cells were recorded along the CA3 transverse axis (Figure 4F), we next assessed whether there was a relation between remapping scores across the full sequence and tetrode location. There was a significant negative correlation between tetrode location and rate difference (*r*_(7)_= −.811, *p*= 0.027) (Figure 4G, Table 4). There was however no correlation between tetrode location and place cell properties (Table 5), spatial correlation (Figure S3B), nor subset switching (Figure S3C). This finding points to greater rate remapping in place cells recorded from the most proximal CA3, compared to distal CA3.

## Discussion

Being raised in a complex environment enhances memory discrimination in rodents^1–6^. Here, we investigated putative neural mechanisms supporting enrichment-dependent memory improvements using a behavioural task that models the automatically encoded and integrative nature of episodic memory. We found that EE-dependent improvements in fine memory discrimination are related to increased levels of adult hippocampal neurogenesis and enhanced sparsity of neuronal activity across the hippocampus. Critically, we also found that EE changes the way CA3 encodes changes in context. EE leads to increased spatial tuning of CA3 place cells, as CA3 place cells in enriched rats carry greater spatial information, have higher firing rates and show increased remapping in response to small geometric alterations to the environment.

These findings are consistent with research showing that adult hippocampal neurogenesis, which is upregulated by enrichment^27^, is fundamental for pattern separation^21^ and stimulation of immature abGCs drives place cell remapping in proximal CA3^34^. The current studies extend this research in two important ways. Firstly, by using a behavioural task that models key features of episodic memory we extend previous findings to the most clinically relevant form of memory. Secondly, we demonstrate enrichment-dependent changes in CA3 place cells, the region that receives input directly from abGCs^22,29^, for the first time. We also show that enriched rodents encode associative memories more efficiently as they require less time to accurately encode novel objects in the context-dependent episodic memory tasks.

Interestingly, the effects of EE on sparsity of activity were particularly evident in the SPDG, compared to the IPDG. Given that the SPDG receives most input from LEC^36^, these findings support recent research suggesting that immature abGCs drive inhibition in response to LEC input^24^, in particular, and increased inhibition in the SPDG, rather than the IPDG, is related to improved pattern separation^37^. Additionally, the finding that EE results in reduced activity in proximal, rather than distal, CA3 points to a special role for proximal CA3 in pattern separation. This is consistent with previous research pointing to proximal CA3 contributing to pattern separation^30–32^, and with our finding of greater rate remapping in cells recorded from proximal, rather than distal, CA3.

The increased spatial tuning of CA3 place cells in the enriched rats suggests that EE leads to more selective and more precise spatial information encoding in CA3, which could support better spatial memory. Combined with recent reports of enrichment-dependent increases of spatial tuning and spatial information in DG mature granule cells in novel environments ^38^, these findings show that EE has clear effects on spatial tuning of place cells in the DG/CA3 network. Interestingly, our results stand in contrast to findings from CA1^28^ which suggests that EE’s effects on spatial tuning of place cells might be limited to DG/CA3.

EE also resulted in increased firing rates in CA3 place cells, consistent with previous reports of enrichment-dependent increases in activity in DG mature granule cells and CA1 place cells^39^. This contrasts with studies showing that increased neurogenesis is typically associated with decreased activity in the DG and CA3^19,22–26^. This discrepancy, however, comes from the way activity is measured. Most studies looking at how EE and adult hippocampal neurogenesis modulate hippocampal activity have focused on differences in percentage of active neurons^28^, rather than on differences in firing rates. The findings of this study suggest that EE reduces the number of active cells, while increasing the activity of those cells. This allows the hippocampus to reduce interference by encoding spatial information using a small neuronal population with high spatial information^38^.

We also report enrichment-dependent increase in rate remapping in CA3 across the different shapes of the morph box. Interestingly, our standard-housed rats did not show CA3 rate remapping in the morph box as previously reported^16,17,33^. This can potentially be explained by slightly different protocols. Here rats had a short exposure to the box during training and morph box shapes were made as similar as possible. Previous studies have used different floors and colours for each shape and extensive pretraining^16,17,33^. This makes the discrimination between shapes in our study much harder, which might explain the lack of rate remapping in our standard-housed rats.

Interestingly, CA3 place cells in the enriched rats showed increased rate remapping across delayed exploration of the same shape. It was reported that exercised mice show increased representational drift in CA1 place cells, i.e., CA1 place cells representation of a stable environment changes more rapidly following exercise^40^. It is possible that increased flexibility in CA3 place cell representations in enriched rats comes at the price of decreased stability of representations. This hypothesis would be consistent with adult hippocampal neurogenesis driving forgetting of previous memories and more flexible encoding of novel information^41–44^. Future studies should further assess the effect of EE on flexibility and stability of hippocampal representations, by looking into the effect of EE on place cell representation of familiar and novel environments over days or weeks. This would allow the investigation of whether and, if so, how EE modulates the balance between flexibility and stability in the hippocampal codes.

Finally, CA3 place cells in the enriched rats showed increased subset switching, i.e., more CA3 place cells turned on and off in response to changes to the environment in the enriched rats compared to the standard-housed rats. Increased subset switching suggests enhanced sparseness of activity in the enriched rats, resulting in more specific spatial representations. The increased spatial tuning in CA3 place cells reported here would also point to more precise encoding of spatial information in the enriched rats.

In summary, we report for the first time that EE changes the way the CA3 place cells encode spatial environments, by increasing their spatial tuning and chances of remapping. Together, these findings point to more precise, and more flexible, memory representations in the CA3 of enriched rats. This increase in spatial tuning and in the ability to respond to alterations to familiar environments might provide a novel neuronal mechanism for enrichment-dependent improvements in fine memory discrimination.

## ACKNOWLEDGEMENTS

We thank the members of the St Andrews Animal Unit (SMAU) for their support and advice on the environmental enrichment used in these experiments. This work was supported by a grant from the BBSRC (BB/X007197/1)

## AUTHOR CONTRIBUTIONS

Conceptualization and methodology, S.V., J.A.A.; Investigation, S.V.; Formal Analysis, S.V.; S.D.; Resources, J.A.A.; Writing – Original Draft, S.V.; Writing – Review and editing, J.A.A., S.V., Supervision, J.A.A.; Funding Acquisition, J.A.A.

## STAR Methodology

### RESOURCE AVAILABILITY

#### Lead contact

Further information and requests for resources and reagents should be directed to and will be fulfilled by the lead contact, James Ainge

#### Materials availability

This study did not generate new unique reagents.

**Data and code availability**

### EXPERIMENTAL MODEL AND SUBJECT DETAILS

#### Animals

34 male Lister Hooded rats (Envigo, UK; average weight: 100-125g) were used in these experiments. 24 rats (EE: 12, ST:12) were used in the behavioural and Fos experiments, 10 (EE: 5, ST:5) were used in the recording experiment. All animals were housed in diurnal light conditions (12h light/dark cycle) with *ad libitum* access to water. Testing occurred in the light phase, 6-7 days a week. To encourage exploration during the tasks, the rats were food restricted to no less than 90% of their free feeding weight (15-25g per day). All experiments and surgeries were conducted under project licenses acquired from the UK home office and in accordance with national (Animal [Scientific Procedures] Act, 1986) and international (European Communities Council Directive of 22 September 2010 (2010/63/EU) legislation governing the maintenance of laboratory animals and their use in scientific research.

### METHODS

#### Housing conditions

Animals in the EE group were housed in groups of 4 (behavioural and Fos experiments) or 3 (recording experiment) in an enriched environment consisting of 3 standard cages connected with tunnels. The enriched environment was equipped with one disc running wheel, two polycarbonate houses, two hanging tubes, wooden ladders, and swings (Top, Figure S1A). These were maintained in the cage but moved around weekly. A new assortment of objects was introduced in the environment weekly, and sunflower seeds, banana chips and monkey peanuts were scattered in the environment twice a week to promote exploration. Animals in the standard group were housed in groups of 3 in standard laboratory cages (Bottom, Figure S1A). Each cage was equipped with a polycarbonate house, a hanging tunnel and nesting materials. For the recording experiment, following tetrode implant surgery, all animals were housed individually in high-top cages. However, to maintain enrichment exposure in the EE group, the EE rats were placed into an enriched environment (play pen) with their cage mates for 1 to 3 hours a day 6 to 7 days a week for the duration of the experiment. To habituate the animals to the play pen, daily exposure to the play pen started one month prior to surgery. The play pen was a squared environment with black wooden walls (wall length: 1 meter) equipped with toys of different materials and dimensions, a climbing wall, several wooden and plastic houses as well as cardboard mazes (Figure S2A). The play pen was in a separate room from the room where place cell recording took place, and exposure to the play pen always occurred after the recording for that day had finished

#### Behavioural Experiment & Fos Experiment

##### Apparatus

To manipulate the extent of memory interference in the tasks, the rats were tested on two versions of the Object-Context (OC) and of the Object-Place-Context (OPC) recognition memory tasks (similar and dissimilar conditions). In the similar condition, the test environment was a “morph box” made up of 32 rectangular pieces of white cable trunking (Electrical Supplies, TLC, UK; 7.5cm wide, 50cm high) that were held together in the inner surface with brown tape and masking tape. The morph box could be shaped as a square (each side: 62 cm) or as a circle (diameter: 79 cm) (Top, Figure S1B). The square was obtained by imposing a 90-degree bent in the wall every 8^th^ rectangular piece, thus resulting in four walls made up of eight pieces in a straight line. The circle was obtained by joining together two rectangular pieces at a time, which were then arranged as to obtain a 16-sided polygon. The square and the circle had the same smooth black wooden floor. In the dissimilar condition, the test environment was a rectangular wooden box (32cm x 25.5cm x 22cm) that could be configured as Context A (smooth black walls and a black and white wire mesh floor) or Context B (wooden walls and a smooth floor with a red, a green, a yellow and a blue quadrant) (Bottom, Figure S1B). The floor and the walls of the environment were cleaned thoroughly between trials with Safe4 disinfectant (Safe Solutions, Ltd.) to remove odour cues. The environment was lit by two lamps, positioned at equal distances from the box.

##### Objects

The objects used in the tasks were household objects of different shapes, colours and material. These were roughly the size of the rat. The objects used during habituation were not reused during testing. During habituation and testing, one object was placed in the upper right quadrant of the box, and one in the upper left quadrant of the box. The objects were secured in place with white fastening tape (Dual Lock, 3M). Objects were cleaned with Safe4 disinfectant (Safe Solutions, Ltd.) before each trial. New sets of objects were used every day, and copies of each object were used for different trials to cover any odours. The objects used for the *c-Fos* experiment were identical copies of a white and blue ceramic mug, roughly the size of the rat.

##### Habituation

The rats were handled for 10 minutes per day during the week prior to testing. Habituation was repeated twice, once before testing in the similar version of the OC task, once before testing in the dissimilar version of the OC task. Before the similar version of the OC task, the contexts used during habituation were the square and circle versions of the morph box. Before the dissimilar version of the OC task, the contexts used during habituation were the dissimilar contexts A and B. On Day 1, the rats explored each context for 10 minutes in their cage groups. On Day 2, the rats explored each context for 5 minutes on their own. On Day 3, the rats explored each context for 5 minutes on their own. However, this time two random objects were placed in the upper quadrants (left and right) of the box.

##### Experimental Timeline

Testing occurred in two phases: the Behavioural Experiment and the *c-Fos* Experiment. The Behavioural Experiment started four months after the rats had been placed in their housing condition. The *c-Fos* Experiment occurred roughly one month following the end of the Behavioural Experiment.

##### Behavioural Experiment

The Behavioural Experiment occurred in 4 stages: similar OC task, dissimilar OC task, similar OPC task, dissimilar OPC task. Each task consisted of two sample phases and one test phase. Each phase lasted three minutes. In-between phases, the rat was placed in a holding cage for roughly one minute as the objects and contexts were cleaned, and the next testing phase was prepared.

A. OC tasks: In the first sample phase, the rat explored two identical copies of a novel object within one context. In the second sample phase, the rat explored two identical copies of a different novel object within the other context. The test phase occurred in one of the two contexts, and the rat explored one copy of the object that was explored in the first sample phase, and one copy of the object that was explored in the second sample phase. Thus, at test one object was in a context in which it had been experienced before (familiar OC configuration) and one object was in a context in which it had not been experienced before (novel OC configuration).

B. OPC task: In the first sample phase, the rat explored two different novel objects within one context. In the second sample phase, the rat explored copies of the same two objects in the opposite context. However, in the second sample phase each object was in the opposite quadrant compared to where it was in the first sample phase. At test, the rat was presented with two identical copies of one of the objects explored during the sample phases within one context. Thus, at test one object was in a context and in a location in which it had been experienced before (familiar OPC configuration), and one was not (novel OPC configuration).

For all tasks, in the similar condition, the contexts used were the square and the circle versions of the morph box. In the dissimilar condition, the contexts used were the dissimilar contexts described above. The OC and OPC stages lasted 4 days each. Each rat received one testing trial per day. For all tasks, the object that was novel at test, the quadrant and the context in which the object in the novel representation was presented, and the order in which the contexts were presented were counterbalanced across rats.

##### *c-Fos* Experiment

The *c-Fos* Experiment occurred one month after the last day of the OPC dissimilar task and lasted three days. The rats were randomly assigned to either the Familiar or the Novel Condition.

A. Familiar Condition (EE n=5, ST n= 6): Over two days, the rats explored two copies of the same novel object within one of the similar, familiar contexts (circle or square) for 5 minutes each day. On the third day, the rat explored two copies of the same object explored in the prior two days in the same context (familiar object-context association) for 5 minutes.

B. Novel Condition (EE n=6, ST n= 6): Over two days, the rats were presented with two copies of the same novel object within one of the similar, familiar contexts (circle or square) for 5 minutes each day. On the third day, the rat explored two copies of the same object explored in the prior two days, this time in the other similar context (novel object-context association) for 5 minutes.

The context that was novel or familiar on the third day was counterbalanced across rats.

##### Behavioural Data Analysis

The behavioural videos were scored offline by the experimenter. A random sample of 25% of all test videos were scored by an independent scorer blind to condition to check for inter-rate reliability. Reliability between scorers was good with an intraclass correlation of 0.876 (2-way mixed model). The amount of time spent by the rat exploring each object was measured for all sample and test phases. Exploration was defined as moments when the animal’s nose was roughly 2 cm within the object and directed towards it. For all test phases, a discrimination ratio was computed to determine whether the rat spent more time exploring the object in the novel configuration, using the following formula:

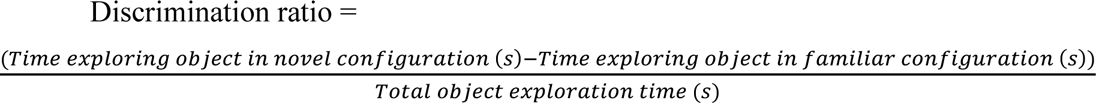

For each animal, average discrimination ratios across the four days of testing were calculated for each task. Average means and standard errors of the mean were then calculated for each group. A positive discrimination ratio indicates that the rat spends more time exploring the object in the novel configuration, signalling memory for the previously encountered object-context (or object-place-context) configuration. Trials were total exploration during sample or test phase was less than 5s were excluded from the analysis. If more than 50% of the trials for a task met exclusion criteria for a rat, data for that rat was excluded from the analysis. This led to the exclusion of 1 ST rat from the Similar and Dissimilar OC and OPC tasks.

##### Histology & DCX quantification

90 minutes following the completion of the test trial of the *c-Fos* Experiment, the animals were given a lethal dose of sodium pentobarbitol and transcranially perfused with phosphate-buffered saline (PBS), followed by roughly 350 ml of 4% paraformaldehyde (PFA). Brains were then extracted and stored in 20% sucrose overnight at 4 degrees. The brains were cut horizontally at 50um on a freezing microtome. For doublecortin (DCX) and *c-Fos* immunohistochemistry, 1 in 8 series were blocked for 2 hours in 20% Normal Goat Serum (NGS) prepared in 0.1% PBS-T (Triton). To stain against DCX, the slices were incubated in a primary antibody solution prepared in 1% NGS in 0.1% PBS-T for 24h at room temperature on a stirrer. The primary antibody was rabbit anti-doublecortin (ABCAM, ab18723, 1:1000). To stain against *c-Fos*, the primary antibody solution contained rabbit anti-Fos (Cell Signalling Technology, 9F6, 1:1000). Slices were washed in PBS 5 times for 2 minutes before being incubated in biotinylated IgG solution (Vectastain Elite ABC Kit, 1:200) for 1h. Slices were the washed in PBS 5 times for 2 minutes before being immersed in avidin-biotin complex (Vectastain Elite ABC Kit, 1:50) for 1h. Sections were then reacted with nickel enhanced 3,3-diaminobenzidine tetrahydrochloride (Sigma) and mounted and cover slipped with DPX. DCX positive cells were imaged using a brightfield microscope (Zeiss ApoTome, 40x). DCX+ cells were quantified throughout the dorsal dentate gyrus (Bregma −3.68mm; Bregma −5.10mm) by an experimenter blind to experimental condition. Cells located in the granule cell layer and the subgranular zone, and whose cell body was clearly visible, were included in the count. For each image, the length of the dentate gyrus was measured in micrometres in ImageJ, and the DCX+ cell density was calculated for each image by dividing the number of DCX+ cells by the length of the dentate gyrus. Average DCX density counts across four dorsal sections were computed for each rat and used in the analysis

##### *c-Fos* Regions of Interest (ROIs)

Regions were identified with reference to the Paxinos & Watson (1991). Counts were taken from 6 subregions of the dorsal hippocampus (Bregma – 3.68, Bregma – 5.10) (suprapyramidal DG, infrapyramidal DG, proximal CA3, distal CA3, proximal CA1, distal CA1).

##### *c-Fos* quantification

Photographs of the regions of interest were taken using a 10x magnification (Zeiss ApoTome). Photographs of at least four, and a maximum of six, sections were taken for each region. Images were processed using ImageJ. To identify *c-Fos* positive cells, ImageJ took a mean grey scale value for each picture. *c-Fos* positive cells were included in the count if their brightness was more than 3 standard deviations greater than the mean grey scale for that picture. Density *c-Fos* counts were obtained by dividing the number of *c-Fos* positive cells by the size of the region they were counted from (in *mm*^2^). To compare across regions with different cell densities, normalized cell density counts were computed by dividing raw cell density counts from each area by the mean count for that area across groups and conditions and multiplying the resulting number by 100.

#### Recording Experiment

##### Surgical implantation of electrodes

Delrin Microdrives (Axona, LtD., UK) contained 8 tetrodes that could be moved independently in bundles of 2. Each tetrode was made of 12.5 micrometre tungsten wire (California Fine Wire, Grover City, CA). Tetrodes were placed in bundles of 2 in 33-gauge x 11 m steel cannulae. Each cannula was glued with superglue (3M, UK) to a plastic shuttle, and each microdrive had 4 plastic shuttles in total. Each shuttle was connected to a screw mechanism. The cannulae were organized in two rows, with 0.4 mm between rows. The cannulae were spaced roughly 0.6mm apart to allow us to target CA3 across the proximodistal axis. This resulted in roughly 1.8 mm between the most far right and the most far left cannulae in the drive. Before implantation, the tetrodes were cut to roughly 5 mm and plated with gold until the impedance of each electrode tip was between 100-200 kW. On the day of surgery, the rats were anaesthetised with Isoflurane in an induction chamber and injected with analgesic Metacam subcutaneously prior to being placed in a stereotaxic frame. The skull was then exposed and the microdrive was implanted in the left hemisphere and targeted at CA3 (3.72-4.0 posterior to Bregma), with the most proximal tetrodes implanted roughly 2.3 mm lateral to midline and the most distal tetrodes implanted roughly 4.1 mm lateral to midline. After the craniotomy, dura was cut, and the tetrodes were lowered 2.8 mm below dura. The implant was the secured to the skull using jewellers’ screws and dental cement. A jeweller screw located near the front of the rat’s skull was connected to the microdrive’s ground wire.

##### Recording

Screening for place cells began one day after surgery. The microdrive was connected to a cable which provided unfiltered electrical signal from single electrodes to the recording system (DaqUSB, Axona Ltd., UK). Within the recording system, the signal was bandpass filtered (600-6000 Hz) and amplified 5000-20000 times with a unity-gain operational amplifier. An oscilloscope on a computer screen connected to the recording system allowed us to visualize filtered signals for each electrode (or channel) of each tetrode. During screening sessions, each channel in the oscilloscope was examined for spiking events. Additionally, population-based EEG signal was examined and sampled at 250Hz. Theta frequency, which is in the range of 8-12 Hz and is characteristic of the hippocampus, was looked for. Light-emitting diodes (LEDs) located on the head stage were used to detect the animal’s location. If a unit was detected, a reference channel where no putative cells were present was used as a reference for the other channels. This was done for each tetrode. Additionally, the gain and the threshold for spiking event detection were adjusted to minimize noise and maximize signal for each tetrode. At the end of a recording session, tetrodes from which putative place cells were recorded were nudged down by roughly 10 micrometres. If no units were detected, the tetrodes were moved down by roughly 60 micrometres.

##### Behavioural apparatus

All recordings occurred in an electrophysiology suite consisting of a towel-line flowerpot (screening location) and a recording location. The recording location consisted of a black-curtained arena. The test box was placed on a black wooden floor positioned on a table in the centre of the arena. A white cue card positioned outside of the testing box and a switched off lamp located within the black-curtained arena functioned as global cues for the environment. The test box consisted of the “morph box” described above. For this experiment, the box was painted matte black to avoid the LEDs on the recording head stage reflecting on the walls, which would have resulted in inaccurate detection of the animal’s location within the box. The morph box could be configured as a square with 62-cm sides, a circle with a 79-cm diameter, or four intermediate polygonal shapes (1:7, 2:6, 3:5, 4:4)^11^. Critically, the centre of each testing environment was always in the same location across the distinct shapes.

##### Testing procedure

In the days prior to surgery, the rats were given 2 10-minute exposures to the square and the circle versions of the morph box for three days. On day 1, the rats explored the environments with their cage mates. On days 2-3, each rat explored the environments on his own. This was done to habituate the rats to the square and circle shapes, and to foraging for pieces of chocolate wheetos randomly thrown in the environment. When putative pyramidal cells were detected in at least one tetrode, testing and recording began. The testing procedure always began with a 10-to-15-minute exploration of either the square or the circle. Each exploration lasted 10-to-15 minutes and consisted in the rat foraging for pieces of chocolate wheetos randomly thrown in the box. The rat was always placed in the box facing the south wall. The recording session consisted of the initial square or circle exploration followed by exploration of the four intermediate shapes in sequential order, exploration of the circle (if the sequence started with the square) or the square (if the sequence started with the circle) and a repetition of the initial shape (full sequence, Top, Figure 4A, Figure S2B). This sequence was counterbalanced across days, as some days the rat experienced the shapes starting with the square, some starting with the circle. In-between exploration of distinct shapes, the rat was placed on the towel-line flowerpot (screening location) for roughly 5 minutes as the environment was morphed and the box was cleaned with Safe4 disinfectant. Data for the full sequence were collected from 3 standard-housed rats and 4 enriched rats. For some of the rats (2 standard-housed and 4 enriched rats), and on some days for the other rats, the session only included some, but not all, of the intermediate shapes (incomplete sequence, Bottom, Figure 4A). Data were included for a rat if the rat explored the first shape, the repetition of the first shape, and at least 2 shapes in-between.

##### Histology

Animals were first anaesthetised with Isoflurane in an induction chamber and placed in the stereotaxic frame. 9V current was then passed through each tetrode to create electrolytic lesions at the tetrode tips, to allow clearer visualization of the tetrode tracks. The animals were then given a lethal dose of sodium pentobarbitol and transcranially perfused with phosphate-buffered saline (PBS), followed by roughly 350 ml of 4% paraformaldehyde (PFA). The brain was then stored within the skull at 4 degrees overnight, until the brain was removed from the skull and stored in 20% sucrose in PBS at 4 degrees for 24h. The brain was then cut coronally on a freezing microtome at 50 micrometres and all sections including the hippocampus were mounted on slides and fixed overnight in a PFA bath. To visualize cell bodies, the brain sections were then stained with cresyl violet, and cover slipped with DPX. The cresyl violet protocol consisted of the slides being submerged in Xylene for 2 minutes, followed by sequential submersion in 100% alcohol and 50% alcohol for 1 minute in each solution. The slides were then washed with tap water for 1 minute and placed in Cresyl Violet for 2 minutes. Prior to being cover slipped with DPX, the slides were then submerged again in water, and in the 50% and 100% alcohol solutions for roughly 1 minute.

##### Tetrode location along the CA3 transverse axis

For each animal the distance between the location of the tetrode tip and the most proximal point of CA3 was measured using ImageJ^32^. This value was then divided by the total length of CA3. This resulted in a value ranging from 0 (most proximal CA3) and 1 (most distal CA3).

##### Place cell identification

TINT (Axona) was used to sort the raw recording data. Spike clusters were first generated using KlustaKwik, which automatically sorts raw spike data into clusters based on principal components (PCs) such as amplitude and energy. The resulting clusters were then either deleted (if they did not look like neuronal spikes) or manually refined to reduce noise. Putative pyramidal cells were distinguished from putative interneurons based on spike width. The recording arena was divided into 5 cm x 5 cm bins. Place fields were constructed by summing the total number of spikes that occurred in each location bin (5×5cm), divided by the time that the rat spent in that location. Place fields were defined as contiguous region of ³ 9 (5cm x 5cm) bins where the minimum firing rate was ³ 0.2 Hz and ³ 20% of the peak firing rate for that cell in that shape. Additionally, for each place field a smoothing value of 5 cm SD Gaussian centred on each bin was applied. Finally, a shuffling procedure was used to define place cells; for all putative place cells, the times at which they fired were shuffled relative to the locations at which they fired. This yielded a randomized distribution of spatial information scores. By doing so, 1,000 randomized rate maps, and spatial information scores, were created for each putative place cell. Any cell with a spatial information score that was above the 99^th^ percentile of this randomized distribution of information scores was considered a place cell^45^. Units with at least one place field in at least one shape within the session in which they were recorded were included in the global remapping analysis. For the rate remapping analysis, units with at least one place field in each shape within the session in which they were recorded were included.

##### Quantification of cluster quality

The quality of each cluster was calculated as previously described^46^ using a MATLAB script which measured a squared Mahalanobis distance for each cluster. In particular, for each cluster *c* with *n* spikes, the squared Mahalanobis distance is calculated as:

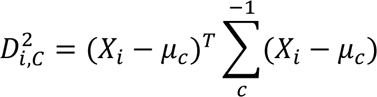

where *X*_*i*_ = any vector with features for spike *I,* and μ_*c*_ = the main vector with features for cluster *c.* Thus, a squared Mahalanobis distance is the distance between the *n*th closest spikes that do not belonging to cluster *c* and the centre of cluster *c.* High values point to better cluster quality and better isolation. Units with an isolation distance of > 20 were classified as highly isolated, units with an isolation distance between 10 and 20 were classified as intermediately isolated and units with an isolation distance < 10 were classified as poorly isolated. Units were included in the analysis if they were intermediately or highly isolated in at least one shape within the session. The automatic isolation distance calculation could only be performed on clusters with a good connection on all four channels per each tetrode. If a channel was grounded or did not have a good connection, isolation distance for that cluster was assessed by comparing those clusters to clusters for which automatic isolation distance could be calculated.

##### Analysis of place cell characteristics

The MATLAB script computed firing maps for each cell, which depict the firing rate for a cell in each bin using a colour ranging from blue (lowest firing rate, cell not active) to red (maximum firing rate, cell maximally active). The firing rate map was analysed for the whole 10-to-15-minute recording trial in each shape, and the following place cell properties were computed: spatial information content, selectivity, spatial coherence, average firing rate in the field, peak firing rate in the field, and place field size. To look at the effect of group on place cell properties, these place cell properties were analysed for all cells that had a place field in the square shape and place cell characteristics in the square were compared for each existing session across groups. The square was picked as a representative shape that could be used to compare the two groups. The spatial information score indicates the amount of information that one can get about the location of the animal based on each spike of the cell. This is calculated as^47^:

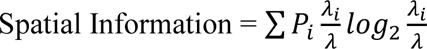

The spatial information score is measured in bits/spike, and it indicates the time that the rat spends in a given bin *i*/total recording time. In the equation, λ_*i*_ = average firing rate in a unit in the *i-*th bin, λ = overall firing rate, and *P*_*i*_ = the probability of finding the rat in the *i* bin. The average in-field firing is calculated by dividing the total number of spikes within the place field by the time the time the rat spent in that location, and the peak in-field firing rate is the maximum firing rate of the cell in its place field (in Hertz (Hz)). Sparsity indicates how specific the place field of a cell is compared to the total area of the test environment. This is calculated as^48^:

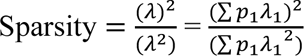

Where λ_*i*_ = the average firing rate of a unit in the *i-*th bin, λ = the overall firing rate, and *P*_*i*_=the probability of finding he rat in the *i* bin. Selectivity indicates how specific the spikes of a cell are to the place field of the cell, and it is calculated as:

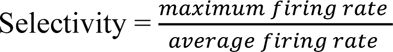

Spatial coherence, instead, is used to determine to what extent firing rates within a pixel are matched with firing rates in adjacent pixels^49^, and it is an indication of how coherent a firing field is. This is calculated as the z-transformed correlation between firing rates in each pixel and the firing rates in eight adjacent pixels of the environment.

Finally, if the same unit was recorded over more than one day, the recording in which the cell had the highest spatial information score was included in the analysis.

##### Analysis of global remapping and subset switching

Place cells that had a field in at least one shape within the session in which they were recorded were included in the main global remapping analysis. This was done using a MATLAB script by correlating rate maps of a cell across the initial shape (either square or circle) and each of the subsequent shapes in the session. Each pixel of the rate map from the first shape in the session was correlated with each pixel of the rate map of the same cell in each of the subsequent shapes within the same session. This generated a Pearson’s correlation coefficient for each comparison (here referred as trial). Correlation values ranged between 0 and 1, with 0 indicating maximum remapping and 1 indicating no remapping. Critically, pixels that were not visited by the animal were discarded from the analysis. Cells that did not have a field in one or more of the shapes in the session, or whose fields were only present in one shape, were considered as subset switching, which is a specific form of global remapping. Finally, for each animal a global remapping value was computed. This was done by calculating, for each animal, the average spatial correlation value for all cells recorded from sessions including the full sequence. Averaged values from each animal were then used in the correlation with the tetrode location values.

##### Analysis of rate remapping

The rate remapping analysis was conducted on place fields, rather than on units. Place fields that were present in all shapes within the session in which they were recorded were included in the rate remapping analysis. Rate remapping was calculated across the first shape in the session and each subsequent shape as the normalized “rate difference” between two^16^. This was done by taking the absolute difference in firing rate for a field between two shapes and dividing it by the sum of the firing rates for that field across the same two shapes. Rate remapping values were calculated for both the average in-field firing rates and the peak in-field firing rates. Greater values indicate greater rate remapping across the two shapes. Finally, for each animal an average rate remapping value was computed. This was done by calculating, for each animal, the average rate difference across the full sequence for all cells, and then averaging that number across all cells recorded for each animal. Only cells that were recorded from sessions including the full sequence were included in the analysis. Averaged values from each animal were then used in the correlation with the tetrode location values.

### QUANTIFICATION AND STATISTICAL ANALYSIS

Statistical analyses were conducted using SPSS (IBM, version 28). Prior to analysis, normality of data was checked using the Shapiro-Wilk test. If the Mauchly’s test was significant, Greenhouse-Geisser corrections were used. To compare rat performance across groups, univariate ANOVAs were run for each task on the discrimination ratios for each group (EE vs ST). To determine whether the average discrimination ratios were significantly greater than chance level, one-sample t-tests against the value of 0 were conducted on the discrimination ratios for each group for each task. To compare object exploration during the test phase across groups, univariate ANOVAs were run for each task on the total object exploration during test (s) for each group (EE vs ST). Additionally, a Mixed Effect ANOVA with Sample (1 vs 2), Task (OC and OPC) and Similarity (Similar vs Dissimilar) as within-subjects factors and Group (EE vs ST) as between-subjects factor was run on the total object exploration (s) to assess differences in the amount of time the rats spent exploring the objects during encoding. A univariate ANOVA was used to compare DCX density counts across the enriched and the standard groups. To determine whether there was a relationship between performance in the tasks and levels of adult hippocampal neurogenesis, Pearson’s product-moment correlation coefficients were calculated with DCX density against discrimination ratios for each task for each animal. *c-Fos* density counts and normalized *c-Fos* density counts were analysed in the dorsal hippocampus (SPDG, IPDG, pCA3, dCA3, pCA1, dCA1). A Mixed Effect ANOVA with Group (Enriched, Standard) and Condition (Familiar, Novel) as between-subjects factors and ROIs as within-subjects factor was used to compare *c-Fos* density counts and normalized *c-Fos* density counts across regions, groups and conditions.

To determine whether there was a significant difference across the enriched and standard groups in place cell properties, Kruskal-Wallis H-tests were conducted with Group (Enriched and Standard) as Fixed Factor. For the place cell properties, the values for the shape “square” were included in the analyses for each cell that had a place field in the square shape. This was done to assess group differences in place cell properties within a representative shape. For the rate remapping analysis, a Generalized Linear Mixed Models (GLMM) were run with Cell as Random Factor and Group and Trial as Fixed Factors on rate differences from place fields that were recorded in sessions that included the initial shape (square or circle), a repetition of the first shape and at least two shapes in-between. This was done to include cells recorded from incomplete sessions. To be included in the rate remapping analysis, the place field had to be present in all shapes within the session. Following significant interactions, Bonferroni-corrected pairwise contrasts were performed to compare rate differences across trials and groups. For the main global remapping analysis, a GLMM was run with Cell as Random Factor and Group and Trial as Fixed Factors on spatial correlation values across the first shape and each subsequent shape. This analysis included all units with a field in at least one shape in the session. To determine whether a higher percentage of place cells showed subset switching across the enriched and the standard groups, observed frequencies of place cells showing subset switching were compared across experimental groups using a Chi Square test of independence. Finally, Pearson’s correlations (or Spearman’s correlation) were run between global remapping scores/rate remapping scores and tetrode location values (distance from the most proximal side of CA3) for each animal.

## SUPPLEMENTAL INFORMATION

**Figure S1: (A)** Example pictures of the enriched environment (Top) and the standard cages (Bottom). (B) Example pictures of the similar contexts (Top) and dissimilar contexts (B) used in the behavioural experiment. **(C)** Relationship between levels of adult hippocampal neurogenesis and performance in the OPC tasks. Scatterplots of the density of DCX+ cells plotted against the discrimination ratios for the Similar OPC (Left) and Dissimilar OPC (Right) tasks. Line of best fit (dashed black) is shown for each plot.

**Figure S2: (A)** Example picture of the play pen**. (B)** Example pictures of the distinct shapes of the morph box (square, 1:7, 2:6, 3:5, 4:4, circle). **(C)** Schematic of the recording procedure involving the incomplete sequences. **(D)** Bar graphs showing the rate differences across the first shape in the recording sequence and each subsequent shape per each animal in the enriched (blue) and standard (orange) groups. Error bars indicate SEM. **(E)** Line graph showing the average spatial correlation scores across the first shape in the sequence and each subsequence shape for cells recorded from the enriched and the standard-housed rats. Cells with a field in all the environments in the recording session were included in this analysis. Error bars indicate SEM.

**Figure S3: (A)** Stacked bar plot comparing the proportion of place cells that showed subset switching (turning on an off) across the recording sequence for each of the enriched (blue) and standard-housed (orange) rats. (B and C) Relationship between tetrode location and spatial correlation (B) or subset switching (C) Scatterplots of the tetrode location plotted against the spatial correlation values (B) or the number of place cells showing subset switching (C) across the sequence. Line of best fit (dashed black) is shown for each plot.

**Supplementary Table S1:** Place fields from enriched and standard housed rats in the square box included in analysis of CA3 spatial tuning

**Supplementary Table S2:** Place cells that had fields in all shapes (full sequence) from enriched and standard housed rats.

**Supplementary Table S3:** Place cells from enriched and standard housed rats with a field in at least one shape included in analysis of CA3 global remapping

**Supplementary Table S4:** Tetrode location and rate difference scores in enriched and standard house rats

**Supplementary Table 5:** Tetrode location and place cell properties in enriched and standard house rats

*In order to meet institutional and research funder open access requirements, any accepted manuscript arising shall be open access under a Creative Commons Attribution (CC BY) reuse licence with zero embargo.”*

