## Supplementary Information for "Increased flexibility of CA3 memory representations following environmental enrichment"

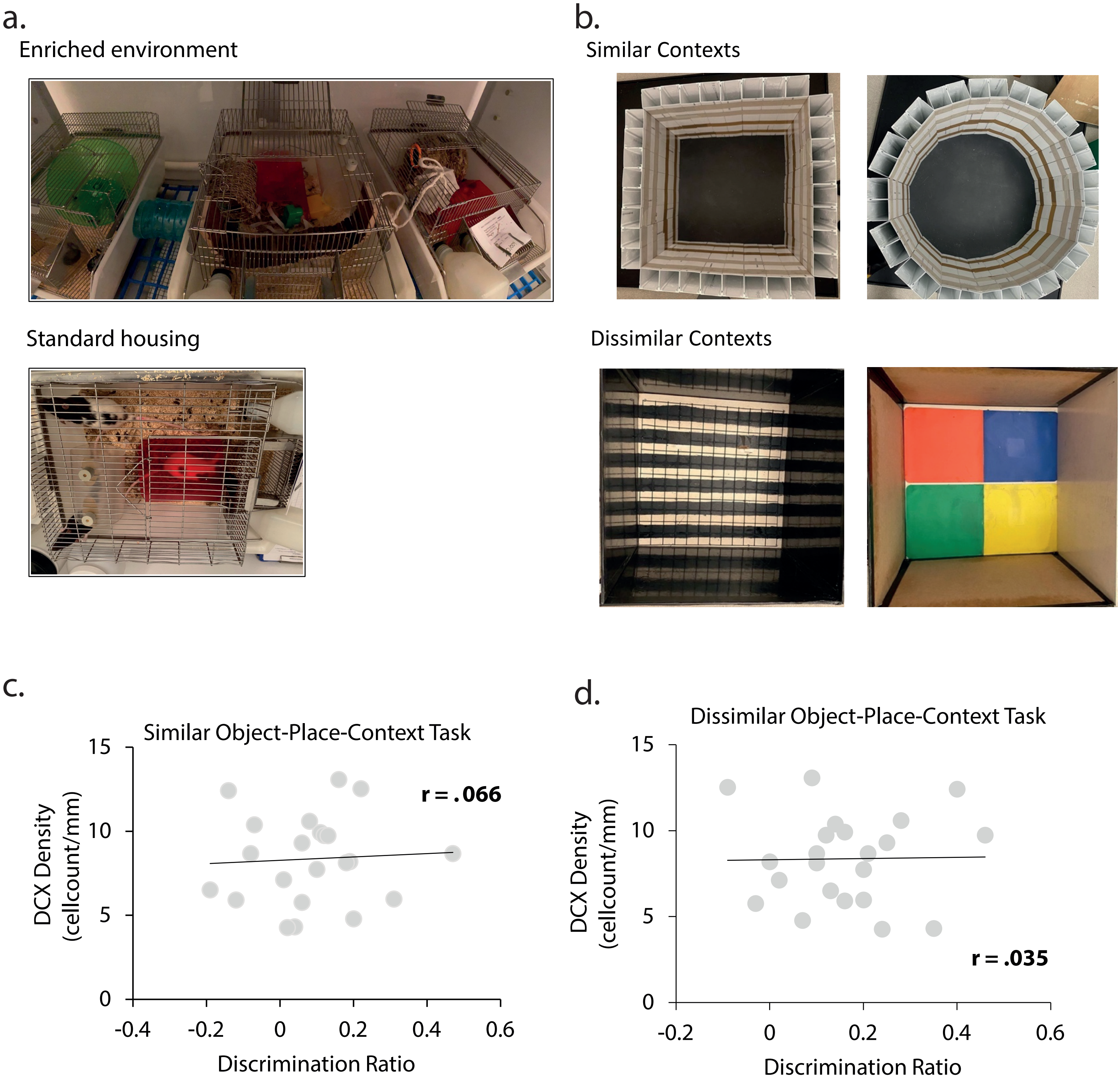
a.
Enriched environment
Standard housing
b.
Similar Contexts
Dissimilar Contexts
c.
Similar Object-Place-Context Task
r = .066
DCX Density
(cellcount/mm)
Discrimination Ratio
d.
Dissimilar Object-Place-Context Task
r = .035
DCX Density
(cellcount/mm)
Discrimination Ratio

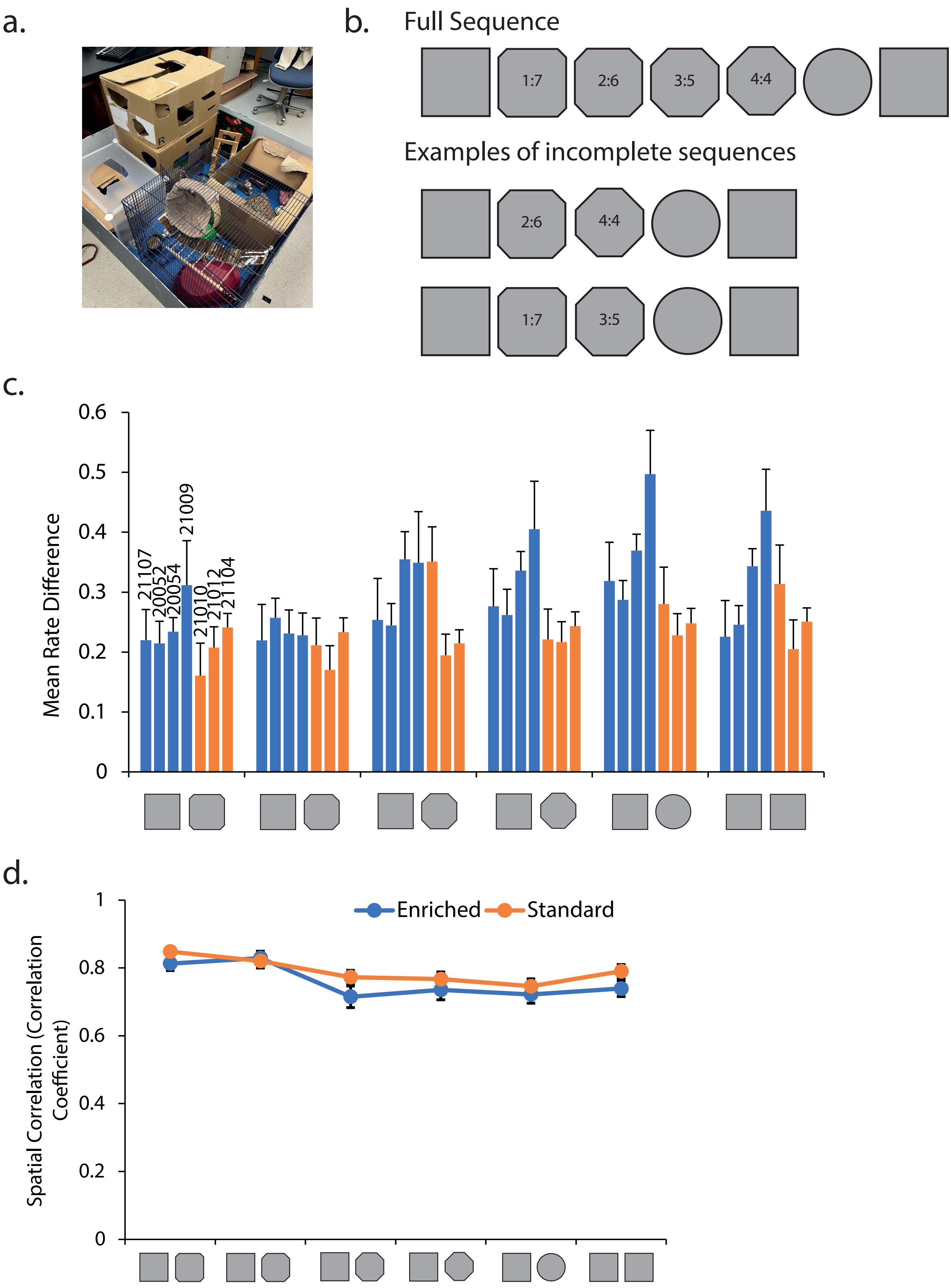
a.
b.
Full Sequence
1:7
2:6
3:5
4:4
Examples of incomplete sequences
2:6
4:4
1:7
3:5
c.
Mean Rate Difference
0.6
0.5
0.4
0.3
0.2
0.1
0
21107
20052
20054
21009
21010
21012
21104
d.
Spatial Correlation (Correlation Coefficient)
1
0.8
0.6
0.4
0.2
0
Enriched
Standard

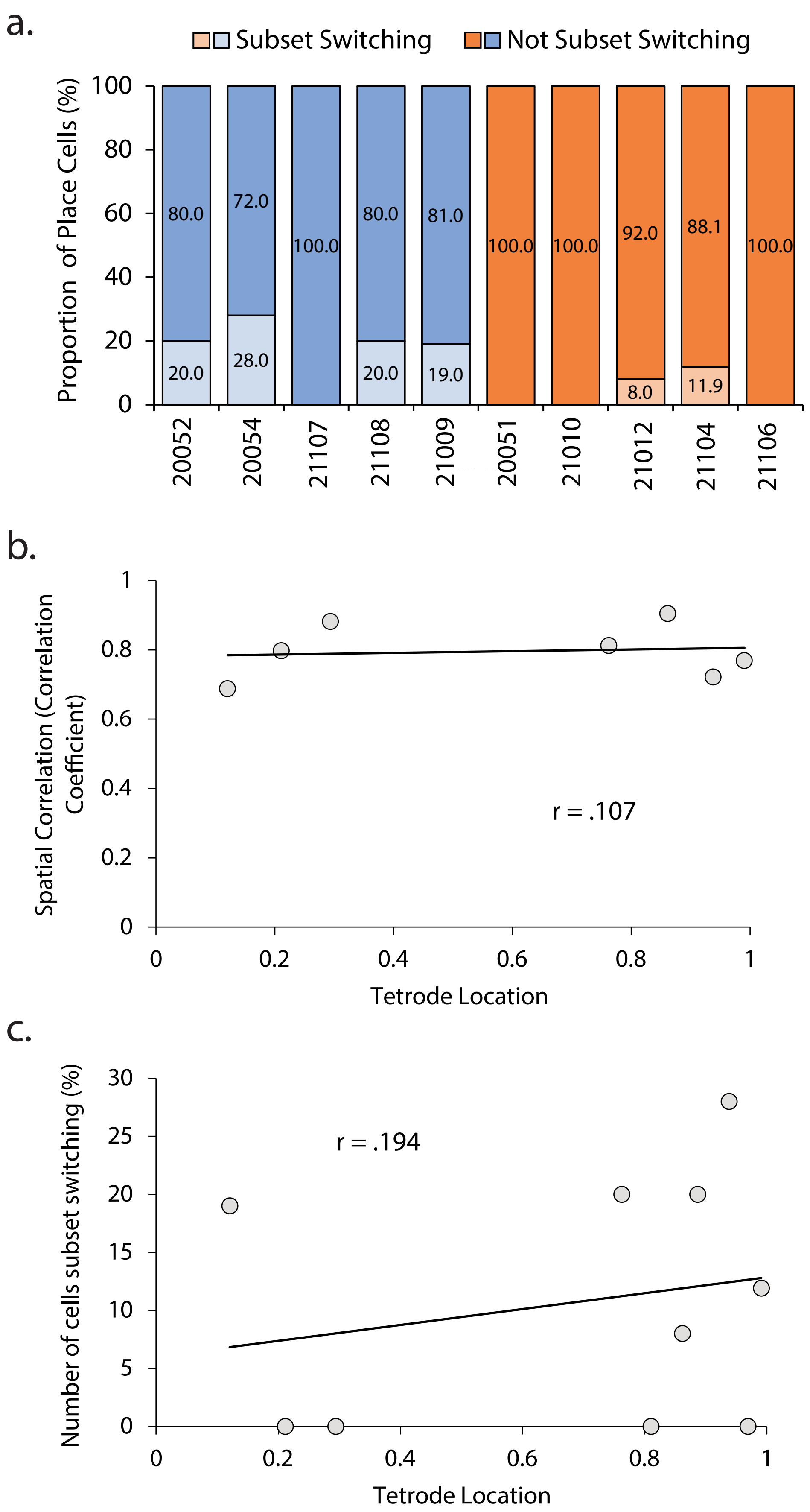
a.
Subset Switching
Not Subset Switching
Proportion of Place Cells (%)
100
80
60
40
20
0
80.0
20.0
72.0
28.0
100.0
80.0
20.0
81.0
19.0
100.0
100.0
92.0
8.0
88.1
11.9
100.0
20052
20054
21107
21108
21009
20051
21010
21012
21104
21106
b.
Spatial Correlation (Correlation Coefficient)
r = .107
Tetrode Location
c.
Number of cells subset switching (%)
r = .194
Tetrode Location

**Supplementary Table 1:** Place fields from enriched and standard housed rats in the square box included in analysis of CA3 spatial tuning

| Rat ID | Group | # Place Fields |
|---|---|---|
| 21107 | EE | 13 |
| 20052 | EE | 46 |
| 20054 | EE | 96 |
| 21009 | EE | 19 |
| 21108 | EE | 5 |
| 21010 | ST | 11 |
| 21012 | ST | 24 |
| 21104 | ST | 108 |
| 21051 | ST | 13 |
| 21106 | ST | 2 |

**Supplementary Table 2:** Place cells that had fields in all shapes (full sequence) from enriched and standard housed rats

| Rat ID | Group | # Place Fields |
|---|---|---|
| 21107 | EE | 10 |
| 20052 | EE | 40 |
| 20054 | EE | 72 |
| 21009 | EE | 18 |
| 21108 | EE | 3 |
| 21010 | ST | 9 |
| 21012 | ST | 22 |
| 21104 | ST | 69 |
| 20051 | ST | 8 |
| 21106 | ST | 2 |

**Supplementary Table 3**: Place cells from enriched and standard housed rats with a field in at least one shape included in analysis of CA3 global remapping

| Rat ID | Group | Place Cells |
|---|---|---|
| 21107 | EE | 13 |
| 20052 | EE | 50 |
| 20054 | EE | 107 |
| 21009 | EE | 21 |
| 21108 | EE | 5 |
| 21010 | ST | 11 |
| 21012 | ST | 25 |
| 21104 | ST | 109 |
| 21051 | ST | 13 |
| 21106 | ST | 2 |

**Supplementary Table 4:** Tetrode location and rate difference scores in enriched and standard house rats

| ID | Tetrode Location | Rate Difference | Group |
|---|---|---|---|
| 21010 | 0.29 | 0.27 | ST |
| 21012 | 0.86 | 0.16 | ST |
| 21104 | 0.99 | 0.22 | ST |
| 20052 | 0.76 | 0.27 | ER |
| 20054 | 0.94 | 0.15 | ER |
| 21009 | 0.12 | 0.31 | ER |
| 21107 | 0.21 | 0.28 | ER |

**Supplementary Table 5:** Tetrode location and place cell properties in enriched and standard house rats

| ID | Tetrode Location | Group | Average Rate | Peak Rate | Bursting | Sparseness | Infoscore | Selectivity | zcoh |
|---|---|---|---|---|---|---|---|---|---|
| 20051 | 0.76 | ST | 1.38 | 2.78 | 0.20 | 0.4 | 0.95 | 9.80 | 3.07 |
| 20052 | 0.94 | ER | 4.15 | 8.05 | 0.18 | 0.26 | 1.59 | 12.96 | 3.27 |
| 20054 | 0.12 | ER | 2.54 | 5.03 | 0.14 | 0.33 | 1.25 | 9.06 | 3.34 |
| 21009 | 0.21 | ER | 3.09 | 5.83 | 0.23 | 0.43 | 0.89 | 6.17 | 3.38 |
| 21107 | 0.89 | ER | 2.77 | 5.53 | 0.16 | 0.55 | 0.56 | 4.51 | 3.24 |
| 21108 | 0.97 | ER | 2.35 | 4.46 | 0.17 | 0.42 | 0.88 | 8.58 | 3.24 |
| 21010 | 0.29 | ST | 1.77 | 3.52 | 0.12 | 0.40 | 1.12 | 8.75 | 3.33 |
| 21012 | 0.86 | ST | 2.30 | 4.72 | 0.12 | 0.50 | 0.74 | 5.61 | 3.62 |
| 21104 | 0.99 | ST | 2.18 | 4.29 | 0.19 | 0.42 | 0.98 | 8.06 | 3.26 |
| 21106 | 0.81 | ST | 1.74 | 3.95 | 0.13 | 0.4 | 0.9 | 5.91 | 3.39 |
